# Metabolic switching promotes microbial coexistence under fluctuating resources

**DOI:** 10.64898/2026.08.19.745668

**Authors:** Yu-Pei Tseng, Jan Engelstädter, Andrew D. Letten

## Abstract

Resource fluctuations can facilitate microbial coexistence when species are sufficiently differentiated in their resource uptake strategies. However, past emphasis on the binary classification of species into equilibrium versus non-equilibrium resource specialists has obscured the potential range of temporal niches available to competitors. Here, we investigate whether microbes that switch between respiratory and fermentative metabolism can coexist with specialist competitors under fluctuations in a single resource. Based on simulations of consumer-resource models, we show that metabolic switching generates distinct growth responses to resource availability, allowing metabolically flexible organisms to exploit temporal variation in ways that differ from specialist strategies. As a result, metabolic switchers can coexist with both respiratory and fermentative specialists under intermediate regimes of resource fluctuation. The competitive ability of metabolic switchers is enhanced when transitions between metabolic states are faster and more responsive to changes in resource availability. Rather than constituting physiological constraints, our results suggest that the metabolic flexibility afforded by overflow metabolism may enable organisms to better exploit fluctuating resources environments.

## Introduction

Natural ecosystems commonly exhibit high species diversity despite seemingly few limiting resources for competitors to partition. Nonequilibrium dynamics are thought to provide one solution to this apparent paradox, whereby resource fluctuations create distinct temporal niches that allow multiple species to coexist (Levins, 1979, Chesson, 1994, 2000). Indeed, recent theory shows that rich opportunities for multi-species (>2) coexistence arise when species have sufficiently differentiated growth responses that allow them to specialise on different properties of a fluctuating resource distribution, including the periodic extremes and asymmetries (Richardson *et al*., 2025). It remains unclear, however, whether the requisite differentiation in species’ growth responses is likely to be commonly found in nature.

The canonical trade-off facilitating coexistence under resource fluctuations is between gleaners and opportunists (Grover, 1997, Yamamichi & Letten, 2022). Gleaner strategists grow efficiently at low resource concentrations and thus dominate under a stable resource supply when the mean resource concentration is kept low. In contrast, opportunist strategists grow faster at high resource concentrations and thus are more competitive when resources are supplied in periodic pulses and variance in resource concentration is high. Provided resource fluctuations are not too extreme, these two strategies can stably coexist via relative nonlinearity of competition – so named because it requires competitors to have intersecting, and therefore differentially nonlinear, per capita growth responses (Grover, 1997, Chesson, 1994, Yamamichi & Letten, 2022). Opportunities for coexistence via relative nonlinearity were long thought to be largely limited to just two species (i.e., a gleaner and an opportunist), but emerging theory shows that the higher moments, beyond variance (e.g. skewness, kurtosis and higher), of a fluctuating resource distribution have the potential to sustain many more coexisting species (Richardson *et al*., 2025).

As with the classic gleaner-opportunist trade-off, specialisation on higher moments necessitates that species per capita growth responses are nonlinear and sufficiently differentiated from each other (Chesson, 1994, Richardson *et al*., 2025). In microbial systems, growth responses are traditionally described by saturating Monod functions (Monod 1949; Holling type II), which exhibit comparatively limited flexibility. While sigmoidal (e.g., Holling type III) growth responses are more flexible and may be sufficient to foster coexistence via moment partitioning, their prevalence in nature remains debated (Kalinkat *et al*., 2023, DeLong *et al*., 2025). Alternatively, phenotypic variation driven by plasticity or rapid evolution has the potential to generate significant variation in the nonlinearity of growth responses (Edwards *et al*., 2013, Yamamichi & Letten, 2021, Letten *et al*., 2024). Nevertheless, to date we have only scratched the surface in explicating the range of plausible pathways by which phenotypic variation can facilitate fluctuation-dependent multispecies coexistence.

One potential mechanism generating the kinds of nonlinear growth responses necessary for moment partitioning in microbial systems is the dynamic switching between metabolic pathways. In carbon catabolism and energy production, microbes primarily rely on respiration or fermentation, which represent alternative strategies differing in rate and efficiency (Pfeiffer *et al*., 2001, Basan *et al*., 2015, Frank, 2010). Respiration generally yields more ATP per unit substrate and yields greater biomass, whereas fermentation provides a lower ATP yield but supports higher metabolic flux and faster growth rates (Bauchop & Elsden, 1960, Pfeiffer *et al*., 2001, Basan *et al*., 2015). These alternative pathways mirror the classic gleaner-opportunist trade-off: respiratory metabolism resembles a gleaner strategy favored under stable, resource-limited conditions, while fermentation corresponds to an opportunistic strategy favored under high-concentration resource pulses. Critically, some microbes are not constrained to one strategy but are able to switch between them via overflow metabolism, shifting from respiration to fermentation at high resource levels even under aerobic conditions (Basan *et al*., 2015). While overflow metabolism is widely studied as a cell-level adaptation to intracellular constraints such as proteome allocation, membrane occupancy, and redox balancing (Basan *et al*., 2015, Zhuang *et al*., 2011, Vazquez & Oltvai, 2016, Vemuri *et al*., 2006), the ecological conditions under which this dynamic flexibility is selectively favored over fixed specialists in natural communities remain poorly understood.

By driving a rate-yield trade-off that shifts cells between gleaner and opportunist dynamics, we hypothesized that overflow metabolism can generate realized functional responses with distinct curvature relative to fixed specialists. In turn, we predicted that these “metabolic switchers” will be able to invade systems comprising resident respirer and fermenter species through partitioning of resource fluctuations, and that they can coexist under appropriate regimes of environmental variability. We investigated this prediction by incorporating metabolic switching into classical consumer-resource models, and analysing competitive dynamics under a range of resource pulsing regimes and model paramterisations.

## Materials and Methods

### Model Formulation

We use a consumer–resource model comprising three microbial species competing for a single limiting resource: two fixed specialists (a respirer and a fermenter) and a metabolic switcher. While each specialist is represented by a single state variable, the switcher is able to switch between two phenotypic states. The general model dynamics are given by:

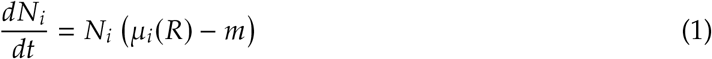

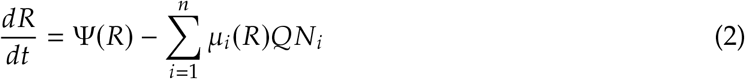

Here, *i* indices consumer identities (four in total: one per specialist, and two for the switcher). Transitions between the switcher’s phenotypic states are governed by a resource-dependent switching function detailed below. *R* is the resource concentration, and *μ_i_*(*R*) represents the per-capita resource-dependent growth rate of consumer *i*. The parameter *m* denotes a mortality rate shared by all consumers, *Q* is the resource quota of a consumer (amount of resource per unit consumer), and Ψ(*R*) describes the resource supply function.

Consumer per capita growth follows a Monod functional response (Monod, 1949):

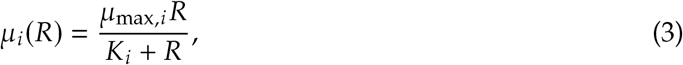

where *μ*_max,*i*_ is the maximum growth rate achievable by consumer *i*, and *K_i_* is the half-saturation constant, i.e., the resource concentration at which growth reaches half of its maximum.

The metabolic switcher differs from other consumers in its ability to shift between two metabolic phenotypes depending on resource availability. Specifically, it can switch between a yield-optimized phenotype (low growth rate but high resource-use efficiency) and a growth-optimized phenotype (high growth rate but low efficiency). To represent this plasticity, we model the switcher as a population composed of two phenotypic states: a yield-optimized state *y* and a growth-optimized state *g*, with resource-dependent switching between them:

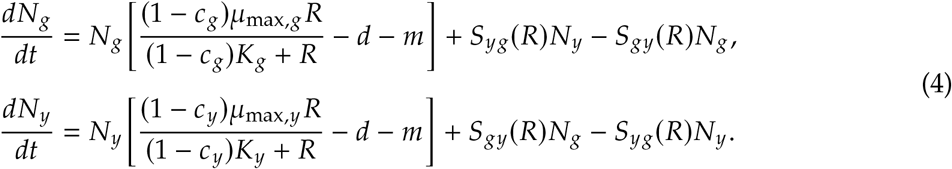

Here, *N_g_* and *N_y_* denote the densities of the growth-optimized and yield-optimized phenotypes of the switching strategy, respectively. To represent the physiological costs associated with maintaining metabolic flexibility, we introduced cost parameters *c_y_* for the yield-optimized state and *c_g_* for the growth-optimized state. This cost was implemented by scaling both the maximum growth rates (*μ_g_* and *μ_y_*) and half-saturation constants (*K_g_* and *K_y_*) of the switcher phenotypes by a factor of (1 ^-^ *c*). This parameterization isolates the metabolic cost to maximum growth rate at high resource concentrations while preserving the initial slope (*α_i_* = *μ*_max,*i*_/*K_s_*_,*i*_), ensuring that low-resource affinity is not penalized. Increasing *c* therefore reduces the competitive performance of the switcher relative to the specialist strategies. When *c* = 0, the switcher incurs no cost and its phenotypes have the same Monod parameters as the corresponding specialist strategies. The functions *S_yg_*(*R*) and *S_gy_*(*R*) represent resource-dependent switching rates from the yield-optimized to the growth-optimized phenotype and vice versa. These switching rates depend on resource availability and are modeled using logistic functions:

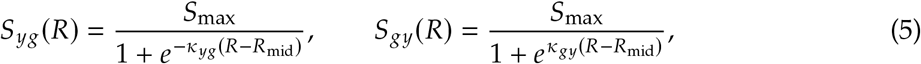

where *κ_yg_* determines how rapidly the yield-optimized phenotype transitions to the growth-optimized state as resource availability increases, whereas *κ_gy_* determines how rapidly the growth-optimized phenotype switches back to the yield-optimized state as resources become limiting. The parameter *R*_mid_ represents the resource concentration at which switching is most sensitive to changes in resource availability (i.e., the inflection point of the logistic function), and *S*_max_ denotes the maximum possible switching rate. Logistic functions were used because they provide a simple and biologically plausible representation of regulatory switching, in which switching rates respond weakly to resource availability when resources are either very low or very high, but respond strongly near a threshold resource level. This captures the idea that metabolic regulation is most responsive near intermediate resource levels, where the relative advantage of fast growth versus efficient resource use changes. The formulation also ensures that switching rates remain non-negative and bounded by *S*_max_ across all resource levels.

Resource supply dynamics differ between continuous and pulsed environments. Under continuous supply, the resource input follows a chemostat formulation:

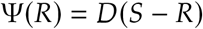

where *D* is the dilution rate and *S* is the resource concentration in the inflowing medium.

In contrast, under pulsed resource supply, the continuous input term Ψ(*R*) is removed from Eq. (2) and replaced by discrete resource additions occurring at fixed time intervals:

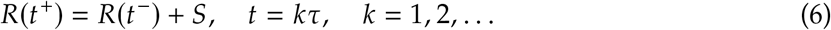

where *τ* denotes the pulse interval, and *R*(*t*^+^) and *R*(*t*^-^) represent the resource concentration immediately after and before a pulse at time *t*, respectively. In this pulsed scenario, *S* corresponds to the size of each resource pulse.

### Model Parameter Selection and Simulation

Model parameters were chosen to represent two contrasting resource-use strategies corresponding to a fermenter and a respirer, analogous to a fast-growing strategist (opportunist) and an efficient slow-growing strategist (gleaner) in classic resource competition theory (Fig. 1A). The fermenter was assigned a higher maximum growth rate and a larger half-saturation constant (*μ*_max,fermenter_ = 0.3 h^-1^, *K*_fermenter_ = 14), representing rapid growth under high resource availability but relatively poor performance at low resource concentrations. In contrast, the respirer was characterized by a lower maximum growth rate and a smaller half-saturation constant (*μ*_max,respirer_ = 0.07 h^-1^, *K*_respirer_ = 0.1), reflecting slower growth but higher resource-use efficiency and competitive ability under resource limitation. For the metabolic switcher, a metabolic cost of *c* = 0.22 was initially assigned, ensuring its growth performance remains slightly subordinate to the specialized respirer and fermenter strategies. This assumption represents a cost of maintaining metabolic flexibility, such that the switcher cannot outperform either specialist under environmental conditions favoring fixed strategies, but may gain an advantage under fluctuating regimes that reward dynamic metabolic adjustment.

**Figure 1.**
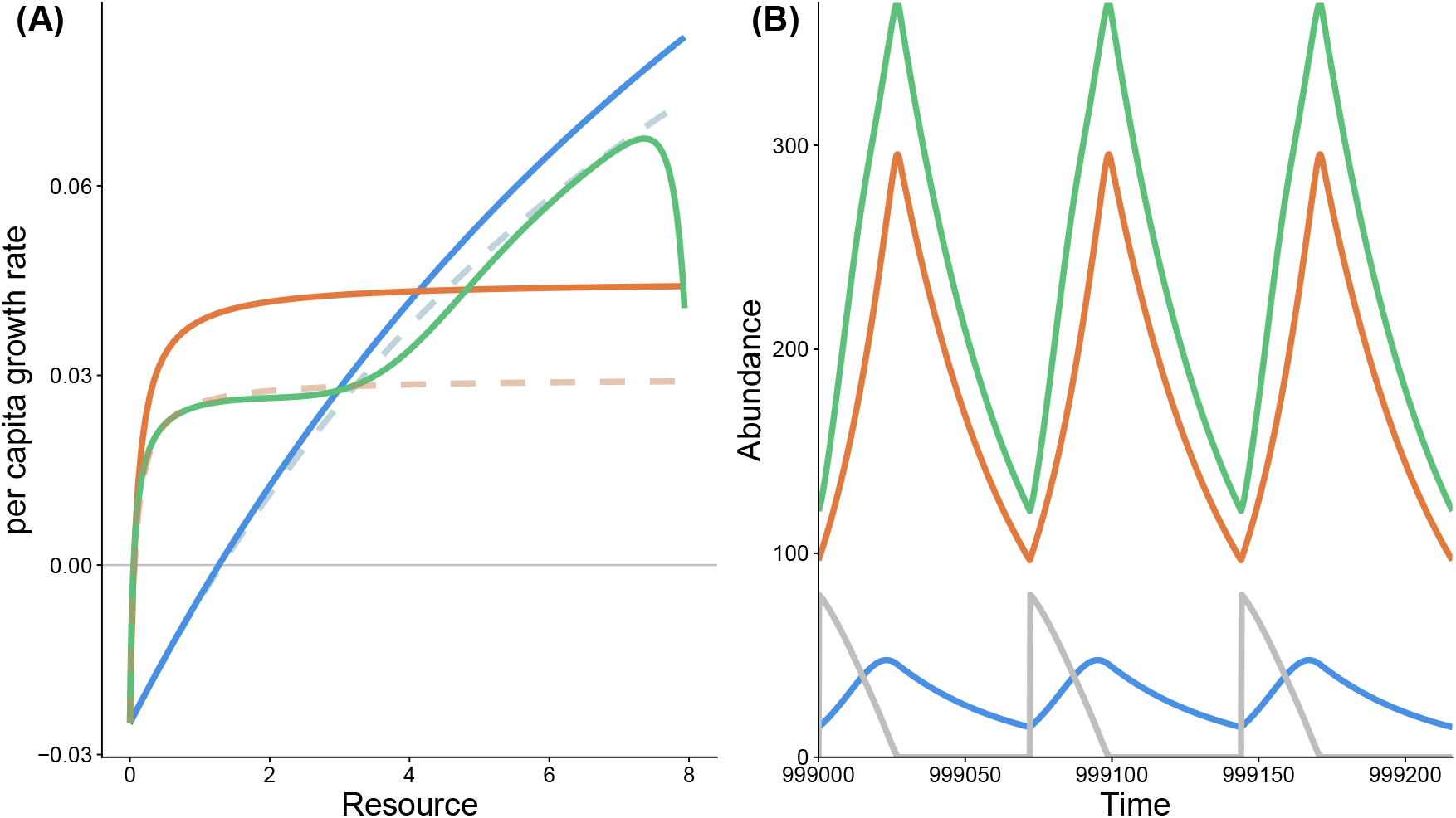
(A) Functional response curves for the fermenter (solid blue), respirer (solid orange), and switcher (solid green) under fluctuating supply of a single limiting resource. The switcher’s per-capita growth rate was quantified in a fluctuating environment as the logarithmic change in total population size, integrating both growth-optimized and yield-optimized states. The intrinsic functional responses of these two states are shown as dashed blue (growth-optimized) and dashed orange (yield-optimized). Due to phenotypic switching, the switcher exhibits a distinct curvature relative to the specialist strategies. Notably, non-monotonic drop in the switcher’s realized response at high resource concentrations arises from a lag in switching back to the growth-optimized state following the arrival of new resource pulses. (B) Population dynamics of the fermenter (solid blue), respirer (solid orange), and switcher (solid green) coexisting under fluctuations in a single limiting resource (grey). For visualization purposes, resource concentration is multiplied by 10 to facilitate comparison with population densities.

Switching was assumed to be symmetric (*k_yg_* = *k_gy_* = 1.4) with a maximum switching rate of *S*_max_ = 1. The midpoint parameter *R_mid_* was set to the resource concentration at which the fermenter and respirer exhibit equal growth rates. For the parameterization used here, this yielded *R_mid_* = 4.1. Consequently, switching is most sensitive near the resource level at which the two specialist strategies perform equally well. For all strategist types, the resource quota and mortality rate were fixed at *Q* = 0.01 and *m* = 0.025.

Initial population densities of the fermenter, respirer, and the yield-optimized state of the switcher were set to 100, whereas the growth-optimized state of the switcher was initialized at a very low abundance at 10^-6^ to ensure solver stability without impacting ecological dynamics. Our system of ODEs was solved numerically over 10^6^ time units and checked for convergence to a periodic steady state. Since the system is driven by external resource pulses, long-term dynamics are expected to approach a stationary periodic pattern rather than a constant equilibrium. To determine whether the system had reached this periodic steady state, we examined population dynamics over the final 10 pulse cycles. Simulations were considered to have reached a periodic steady state when the cycle-end densities of all populations showed no monotonic increase or decrease across these 10 cycles. If the system had not reached a periodic steady state, simulations were extended to 4 × 10^6^ time steps to ensure that transient dynamics had dissipated. Competitive outcomes were determined from the final 10 pulse cycles. For each population, long-term population size was calculated as the time-averaged population density across all time points within these final cycles. A species was considered extinct when its long-term mean abundance fell below 10^-8^ times the maximum abundance attained by any species in that simulation.

We calculated the realized per-capita growth rate of the metabolic switcher in fluctuating resource environments by averaging the logarithmic change in total switcher population size (i.e., *N_y_* + *N_g_*) over fixed evaluation intervals Δ*t* = 1 spanning a complete resource pulse cycle once the system reached periodic steady state. This realized growth rate reflects the combined effects of the functional responses of the yield-optimized and growth-optimized phenotypes, the switching dynamics between phenotypic states, and the frequency and magnitude of resource pulses.

To investigate the effects of metabolic flexibility costs on coexistence, the cost parameter *c* was varied from 0.14 to 0.34 in increments of 0.02. To assess how switching dynamics influence invasion success, we varied the switching parameters controlling transitions between phenotypes, specifically the maximum switching rate (*S*_max_) and the switching slope (*k*). Both parameters were varied from 1.0 to 3.0 in increments of 0.2. Finally, to explore how environmental conditions influence the performance of the metabolic switcher, we varied the size of each resource pulse (*S*) from 7 to 20 in increments of 1, and the pulse interval (*τ*) from 12 to 108 in increments of 12. In addition to the three species competition simulations, we also simulated all three pairwise combinations (i.e, respirer vs fermenter, respirer vs switcher, and fermenter vs switcher) to investigate the frequency of emergent 3-species coexistence. All simulations were conducted using the deSolve package v1.42 in R (Soetaert *et al*., 2010).

## Results

By design, the Monod functional responses (Eq. 3) of the two specialist strategies exhibit the classic gleaner-opportunist trade-off. The fermenter achieves a higher per-capita growth rate at high resource concentrations due to its high maximum growth rate (*μ*_max,*i*_) but performs poorly at low resource concentrations because of its relatively high half-saturation constant (*K_i_*). In contrast, the respirer maintains higher growth rates under resource-limited conditions owing to its lower half-saturation constant, but is outperformed by the fermenter when resources are abundant. The switcher exhibits a realised per-capita growth response that differs markedly from either specialist strategy (Fig. 1A). Unlike the Monod curves of the fermenter and respirer, this response is not an intrinsic functional response, but rather emerges from the interaction between resource dynamics and resource-dependent switching between metabolic states. Consequently, the realised growth response reflects both resource availability and the competitive environment experienced by the switcher. Across the resource gradient, the switcher achieves growth rates intermediate between those of the respirer and fermenter at both low and high resource concentrations, while exhibiting the lowest growth rates at intermediate resource levels near the switching threshold.

Simulations of the full model show that the switcher can stably coexist with both specialist strategies under fluctuations in a single limiting resource (Fig. 1B). This coexistence demonstrates that metabolic switching can generate an emergent realised growth response to resource availability that is qualitatively different from the intrinsic Monod responses of the specialist strategies. By systematically varying the costs associated with the switcher’s yield-optimized and growth-optimized states, we nevertheless find that three-species coexistence occurred within a comparatively restricted region of parameter space (Fig. 2 & S1). This region exhibited a diagonal structure, indicating that coexistence depended primarily on the balance between performance in the two metabolic states rather than on the absolute performance of either state alone. Reductions in performance of the growth-optimized state could be compensated by improved performance of the yield-optimized state, and vice versa, suggesting that successful switching requires maintaining sufficient competitive ability across both resource-rich and resource-poor conditions. Under regimes of larger pulse magnitude and longer inter-pulse intervals, the switcher tolerated reduced performance in both metabolic states while still coexisting with the respirer and fermenter, indicating that these fluctuation regimes increase the fitness advantage conferred by metabolic flexibility. These fluctuation regimes generate prolonged periods of both resource abundance and resource scarcity, increasing the advantage of metabolic flexibility. As a result, the switcher can exploit a wider range of resource conditions and persist despite lower performance in either individual metabolic state. In this regime, the switcher coexisted with the respirer and fermenter despite reduced performance in each state. In contrast to this consistent shift in the 3-species coexistence region, the parameter space corresponding to switcher dominance initially increases with the length of the pulsing interval before contracting again.

**Figure 2.**
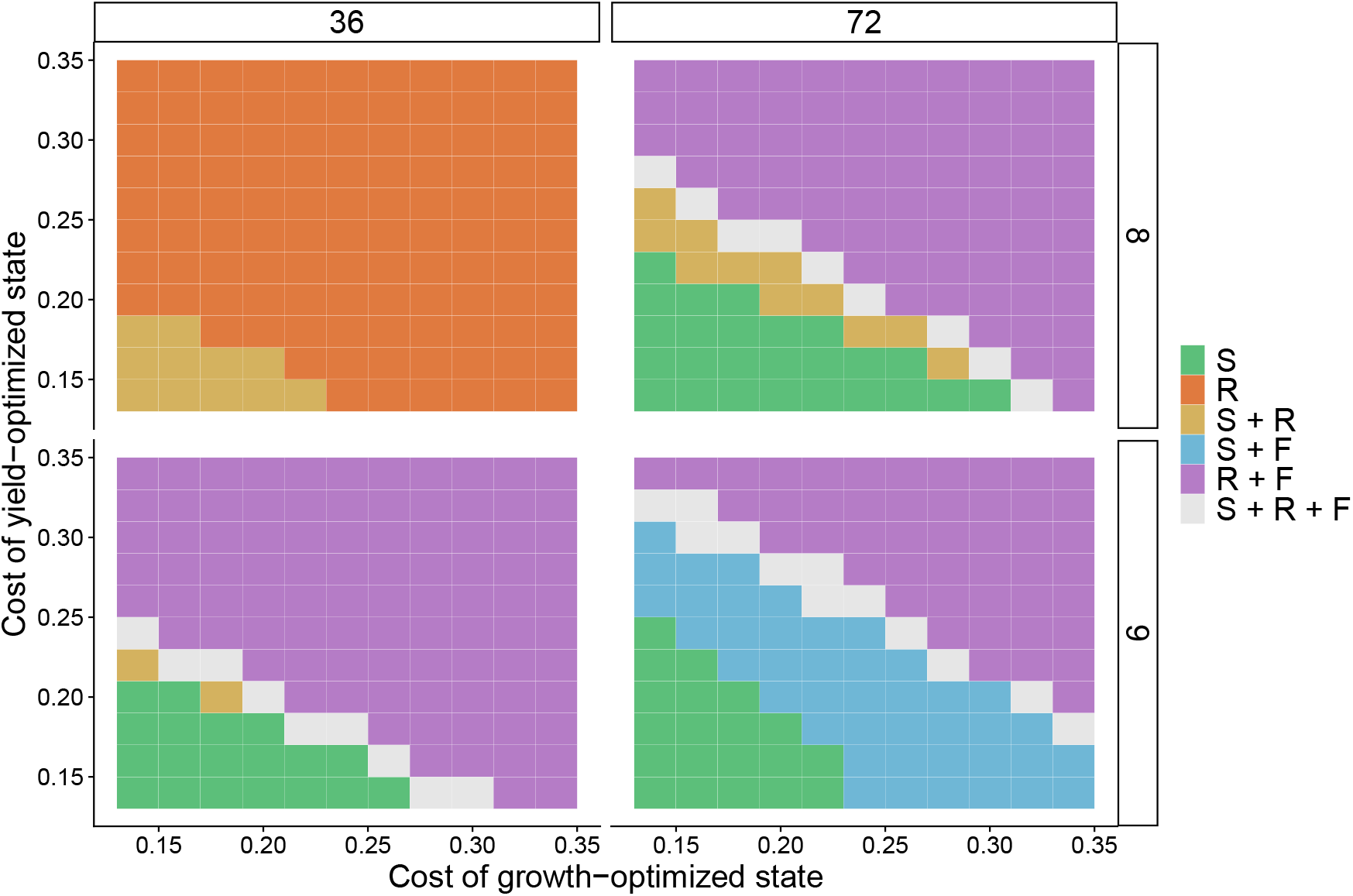
Competition outcomes among the fermenter (F), respirer (R), and switcher (S) across a range of costs and different resource fluctuation regimes. The cost is defined as the reduction of both the maximum growth rate and the half-saturation constant for both the growth-optimized state and the yield-optimized state. Higher costs render the switcher competitively inferior to either specialist. Columns represent different pulse intervals (inverse of resource supply frequency), and rows represent different pulse sizes (resource input magnitude). The region of three-species coexistence (grey) exhibits a diagonal structure, consistent with a trade-off between performance in the switcher’s two metabolic states. Under specific fluctuation regimes—characterized by lower pulse frequency (longer intervals) and higher resource supply—the coexistence region shifts toward higher values of both pulse interval and pulse size. See Fig. S1 for the full resource fluctuation regimes.

Across the fluctuation gradient, distinct community outcomes emerged. Under mild fluctuations (small pulse size and short intervals), both the fermenter and switcher were driven extinct, with the respirer excluding both (Fig. 2 & S1). With increasing fluctuation strength, both the fermenter and switcher were able to persist alongside the respirer. Under intermediate fluctuation regimes, the parameter space supporting switcher dominance reached its maximum extent, while the respirer was progressively excluded. Under stronger fluctuations (infrequent pulses and large pulse sizes), the fermenter increasingly dominated, and the switcher was excluded.

We further examined the effect of switching dynamics by assigning symmetric costs to the switcher’s two metabolic states. Both the slope of the switching function (Fig. 3 & Fig. S2) and the maximum switching rate (Fig. 3, Fig. S3) influenced coexistence outcomes. Increasing the slope and the maximum switching rate of the switching function expands the parameter space over which the switcher persists, enabling its maintenance even at relatively high switching costs. This is intuitive because switching is always biased to the currently optimal phenotypic state in our model, so that faster switching is beneficial.

**Figure 3.**
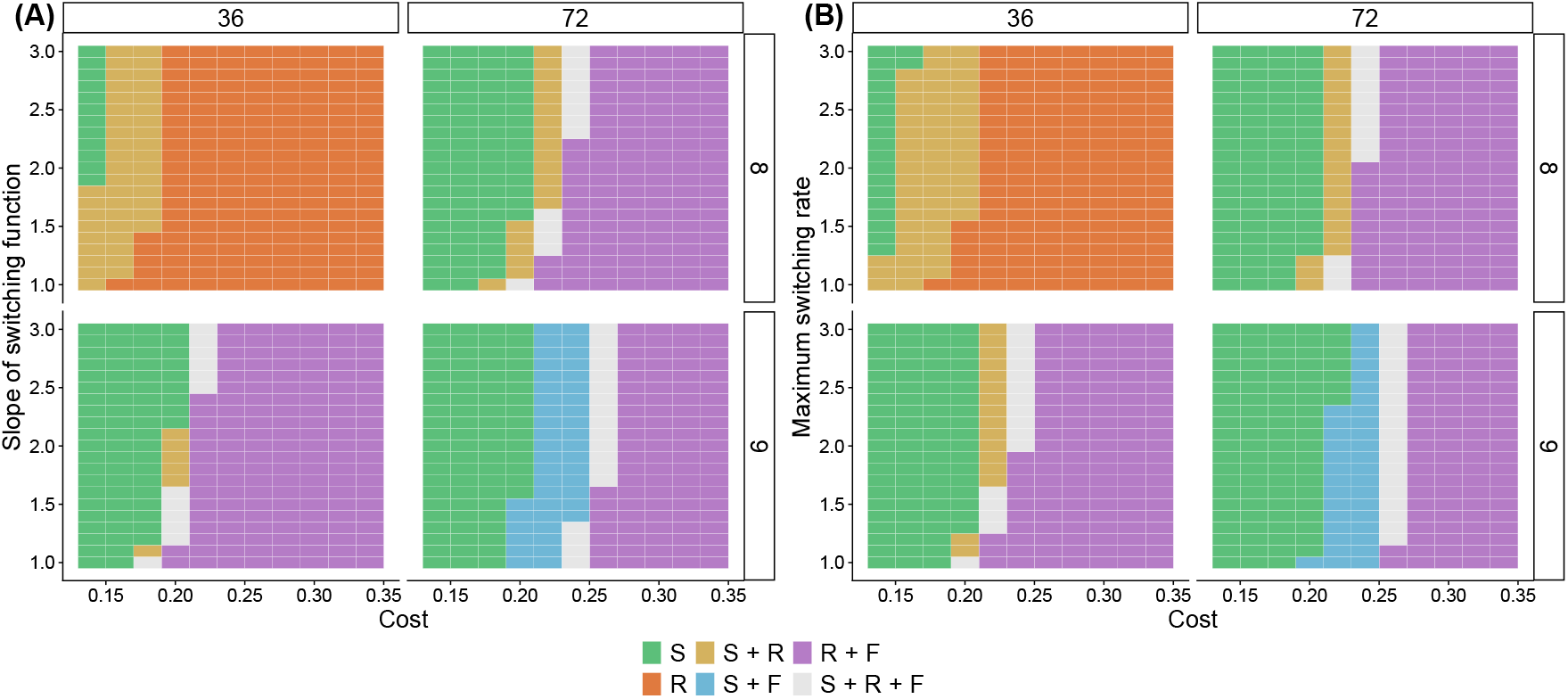
Competition outcomes among the fermenter (F), respirer (R), and switcher (S) across resource fluctuation regimes. (A) Effects of switching cost (assumed symmetrical between metabolic states) and switching-function slope. (B) Effects of switching cost and maximum switching rate. Columns represent different pulse intervals (inverse of resource supply frequency), and rows represent different pulse sizes (resource input magnitude). Increasing the slope of the switching function expands the parameter space over which the switcher persists, enabling its maintenance even at relatively high switching costs. See Fig. S2 for the full resource fluctuation regimes.

Finally, to assess emergent coexistence, we compared the three-species dynamics against those observed for each of the constituent pairs (Fig. S4–S10). In the two-species system (respirer and switcher), the switcher excluded the respirer under conditions with pulse interval = 60– 72 and pulse size = 11 (Fig. S4–S6). However, in the presence of the fermenter, the respirer persisted, resulting in either three-species coexistence or coexistence between the respirer and fermenter (Fig. S1–S3). This indicates that the fermenter modifies competitive dynamics in a manner that permits persistence of the respirer despite pairwise exclusion by the switcher, implying that coexistence in the three-species community cannot be predicted from pairwise interactions alone.

## Discussion

Our results demonstrate that metabolic switchers can coexist alongside specialist strategists in fluctuating resource environments. This observation contributes to a growing body of research demonstrating the wider applicability of the canonical gleaner-opportunist trade-off (Grover, 1997, Yamamichi & Letten, 2021, 2022, Letten *et al*., 2024), while also highlighting a novel pathway to multispecies coexistence via moment partitioning (Richardson *et al*., 2025). More specifically, switching between metabolic states results in distinct and highly nonlinear realised functional responses under resource fluctuations, which allows switcher strategists to better exploit resource pulses of intermediate amplitude and/or frequency than specialist (fixed) strategists. Respirers (~gleaners) and fermenters (~opportunists) in turn dominate under more stable and extreme conditions, respectively.

The temporal dynamics of resource availability are as much a function of the biotic feedbacks that arise from resource consumption as they are of abiotic resource supply (Chesson, 1994, Richardson *et al*., 2025). It is perhaps then unsurprising that we observe emergent coexistence in the respirer-fermenter-switcher system, where the outcome of competition cannot always be predicted from pairwise interactions alone (Friedman *et al*., 2017, Chang *et al*., 2023). For example, all three species were found to stably coexist under conditions where the switcher would otherwise exclude the respirer in the absence of the fermenter. This shift arises because rapid exploitation of high-resource pulses by the fermenter reduces the switcher’s advantage, while simultaneously extending low-resource conditions that favour the respirer.

The extent to which metabolic switching reflects an effective competitive strategy will also be governed by physiological constraints on how rapidly organisms can adjust their metabolic state. For example, a higher maximum switching rate and a steeper slope of the switching function can compensate for a higher metabolic cost of switching. The maximum switching rate represents the upper limit on how rapidly a cell can reallocate metabolic resources between fermentative and respiratory pathways. This rate is constrained by physiological limits (Frank, 2010) such as enzyme synthesis capacity, proteome allocation (Basan *et al*., 2015), and membrane occupancy (Zhuang *et al*., 2011, Vazquez & Oltvai, 2016), which together bound how quickly metabolic fluxes can be reconfigured. These constraints reflect fundamental physical limits on cellular resource allocation and have been proposed as key drivers of overflow metabolism. In contrast, the slope of the switching function represents how rapidly a microbial cell reallocates metabolic fluxes between growth-optimized and yield-optimized pathways in response to resource availability. A steep slope indicates the organism responds quickly to changes in resources, whereas a shallow slope indicates slower adaptation. In real microbial systems, this is akin to the lac operon induction in *Escherichia coli* (Wanner *et al*., 1978) or catabolite repression in *Saccharomyces cerevisiae* (Gancedo, 1998), where gene expression switches sharply once a threshold is crossed. Faster and sharper switching enhances performance in fluctuating environments, but may reduce fitness under stable conditions, where frequent or premature transitions could limit the time spent in the optimal metabolic state (Acar *et al*., 2008).

Our results suggest that temporal variability in resource availability may provide an under-appreciated ecological context in which overflow metabolism is selectively advantageous. While intracellular constraints—such as proteome allocation, membrane crowding, and redox balance (Zhuang *et al*., 2011, Basan *et al*., 2015, Vazquez & Oltvai, 2016, Vemuri *et al*., 2006)—can explain cell-level bioenergetic trade-offs, they cannot explain why metabolically flexible strategies persist alongside fixed specialists in nature. We demonstrate that environmental fluctuations create ecological windows (intermediate frequency and amplitude regimes) where the fitness benefits of switching offset the physiological overheads. It follows that fixed fermentative strategies may also reflect an adaptation, or exaptation, to competition in highly pulsed resource environments.

While we found multiple combinations of different ‘costs of switching’ compatible with three species coexistence, we note that holding one cost fixed (e.g. maximum growth) significantly narrows the parameter space for multispecies coexistence. From this we can infer that complementary mechanisms will likely be important in natural systems. Most notably, by selecting for fermentation and overflow metabolism, fluctuating resource environments may already further promote the maintenance of diversity via metabolic cross-feeding (Carlson *et al*., 2018, Kehe *et al*., 2021). This phenomenon is well illustrated in *Escherichia coli*, where glucose consumption leads to acetate and glycerol secretion (Treves *et al*., 1998, Voegele *et al*., 1993, Millard *et al*., 2021), as well as in *Saccharomyces cerevisiae*, which produces ethanol during fermentative growth (Pasteur, 1857, Gancedo, 1998). Such interactions can promote coexistence by expanding the effective resource space and introducing additional temporal and metabolic niches, thereby complementing fluctuation-driven coexistence mechanisms (Pfeiffer & Bonhoeffer, 2004, MacLean & Gudelj, 2006).

In summary, metabolic flexibility is not universally advantageous, but is expected to be selectively favored under certain temporal regimes of resource supply. These results provide a potential explanation for the persistence of metabolically flexible strategies in natural systems, where the costs of metabolic flexibility are balanced by the benefits of exploiting transient resource pulses. They also raise the possibility that resource pulsing may favor fixed fermentative strategies under conditions where rapid substrate exploitation outweighs reduced energetic efficiency. Finally, these findings generate testable hypotheses on the effects of resource-pulsing regimes on competitive outcomes and evolution in communities comprising a mix of metabolic strategies.

## Supplementary information

**Figure S1.**
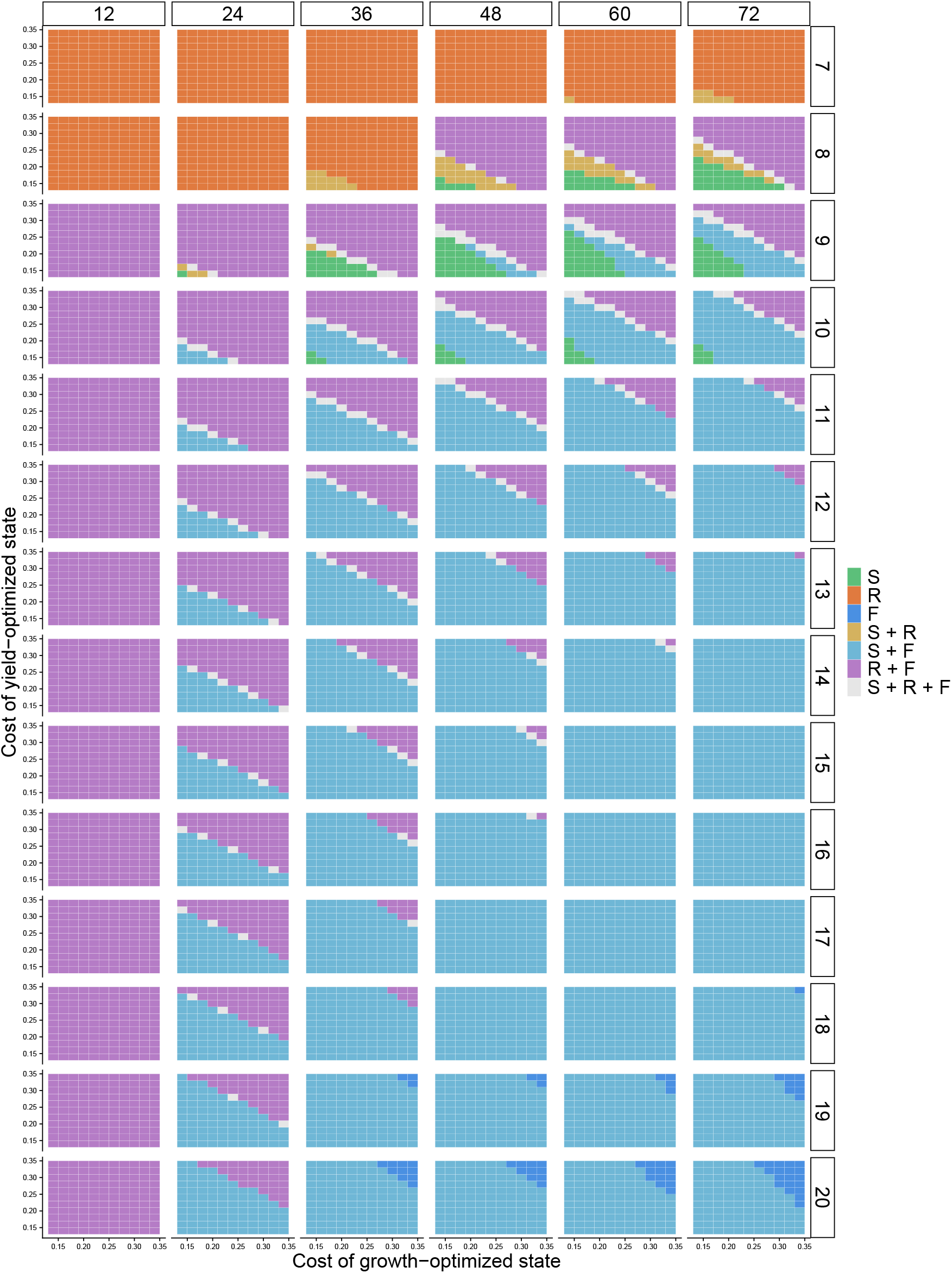
Competition outcomes among the fermenter (F), respirer (R), and switcher (S) across a range of costs and resource fluctuation regimes. Columns represent different pulse intervals (inverse of resource supply frequency), and rows represent different pulse sizes (resource input magnitude).

**Figure S2.**
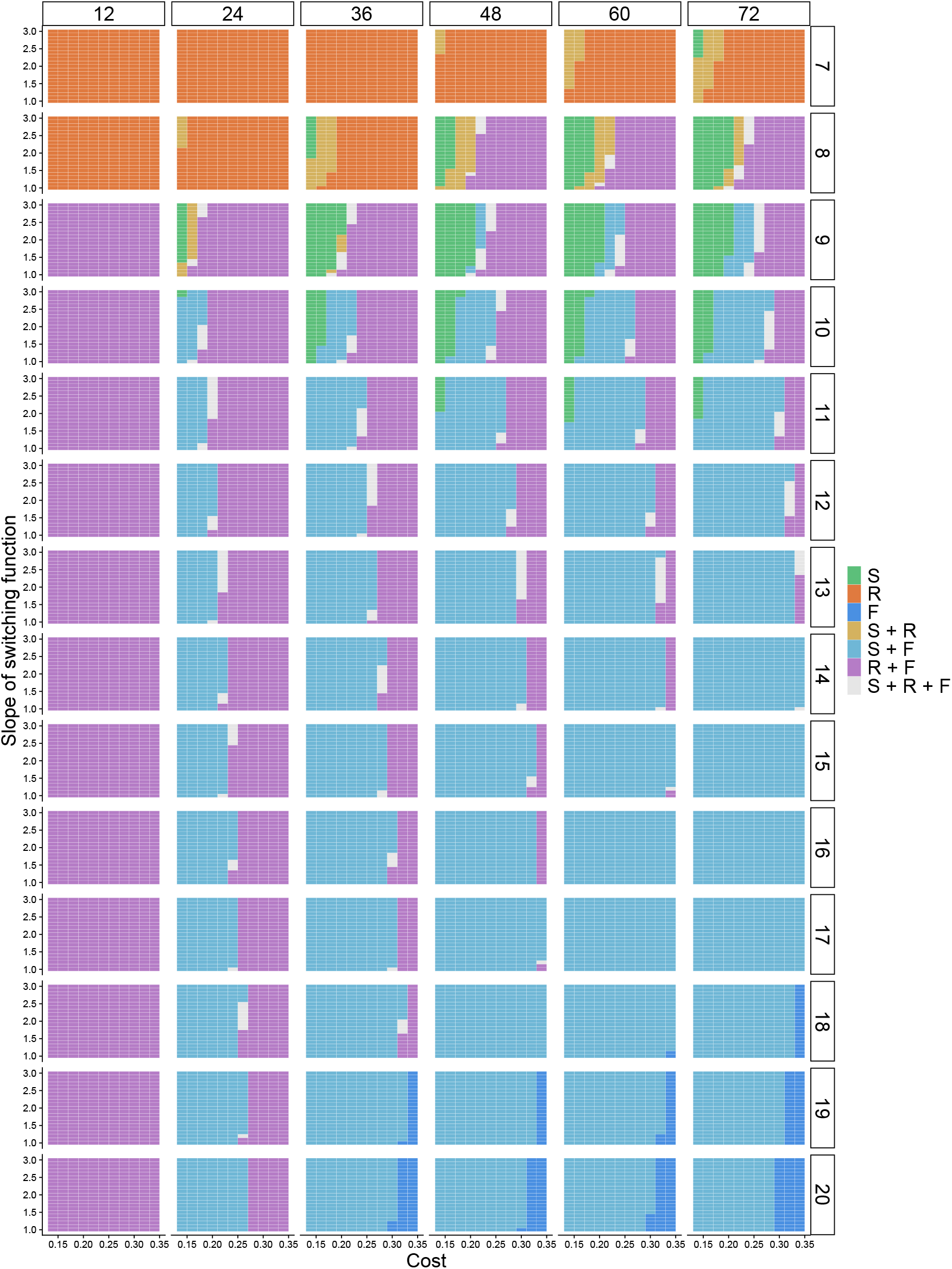
Competition outcomes among the fermenter (F), respirer (R), and switcher (S) across a range of costs (assumed symmetrical between metabolic states), switching function slopes, and resource fluctuation regimes. Columns represent different pulse intervals (inverse of resource supply frequency), and rows represent different pulse sizes (resource input magnitude).

**Figure S3.**
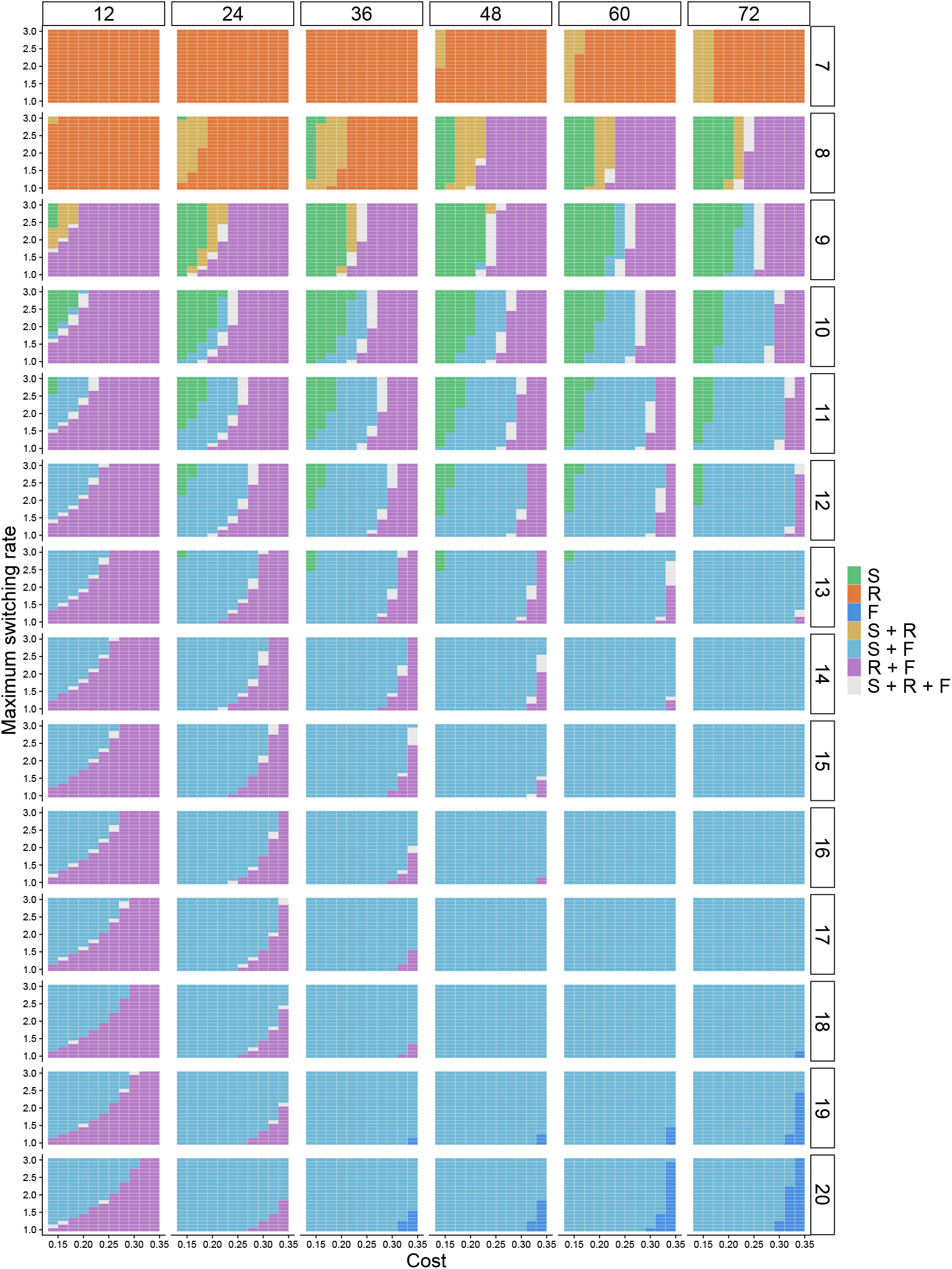
Competition outcomes among the fermenter (F), respirer (R), and switcher (S) across a range of costs (assumed symmetrical between metabolic states), the maximum switching rate of the switching function, and resource fluctuation regimes. Columns represent different pulse intervals (inverse of resource supply frequency), and rows represent different pulse sizes (resource input magnitude).

**Figure S4.**
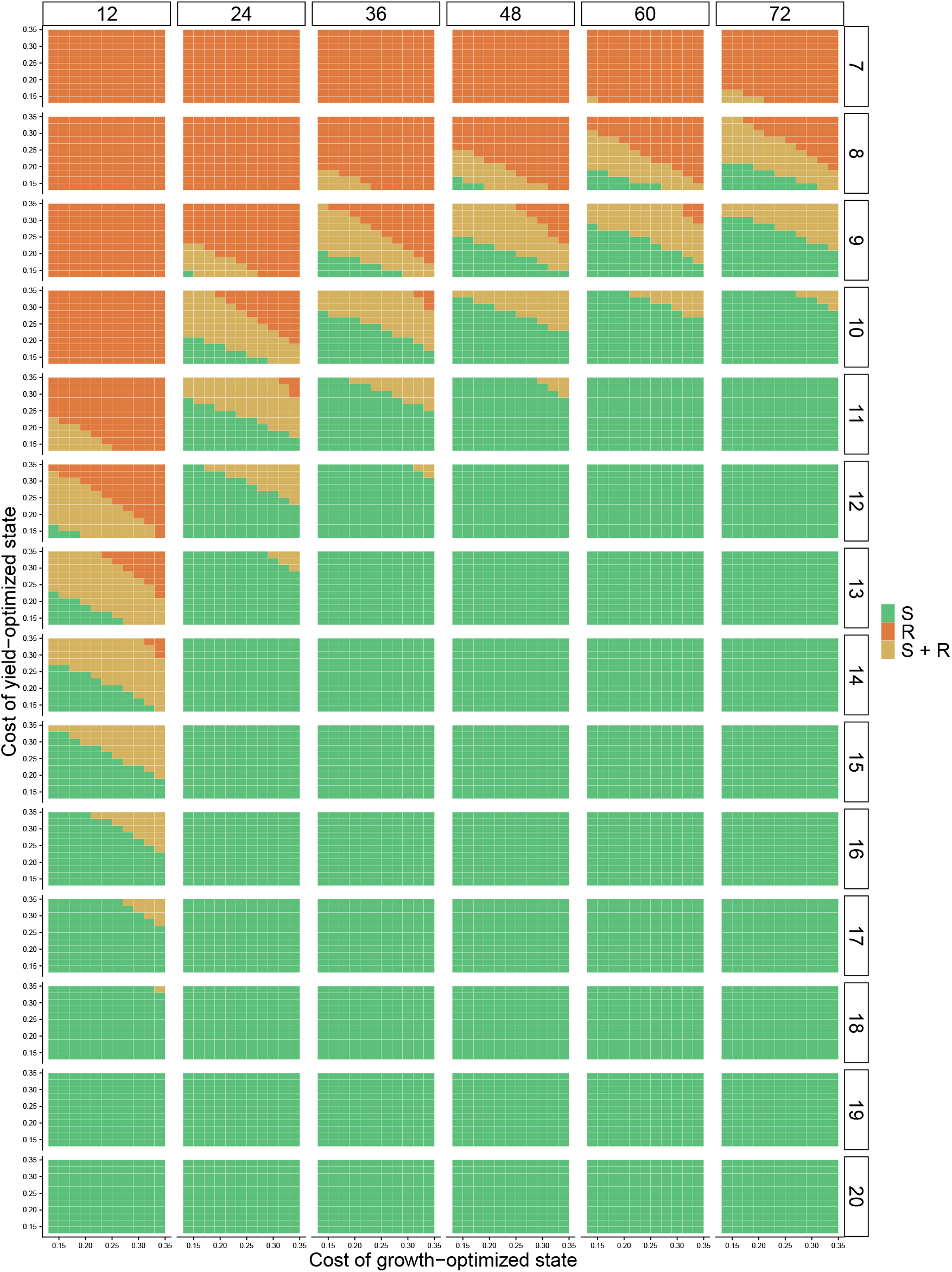
Competition outcomes among the respirer (R) and switcher (S) across a range of costs and resource fluctuation regimes. Columns represent different pulse intervals (inverse of resource supply frequency), and rows represent different pulse sizes (resource input magnitude).

**Figure S5.**
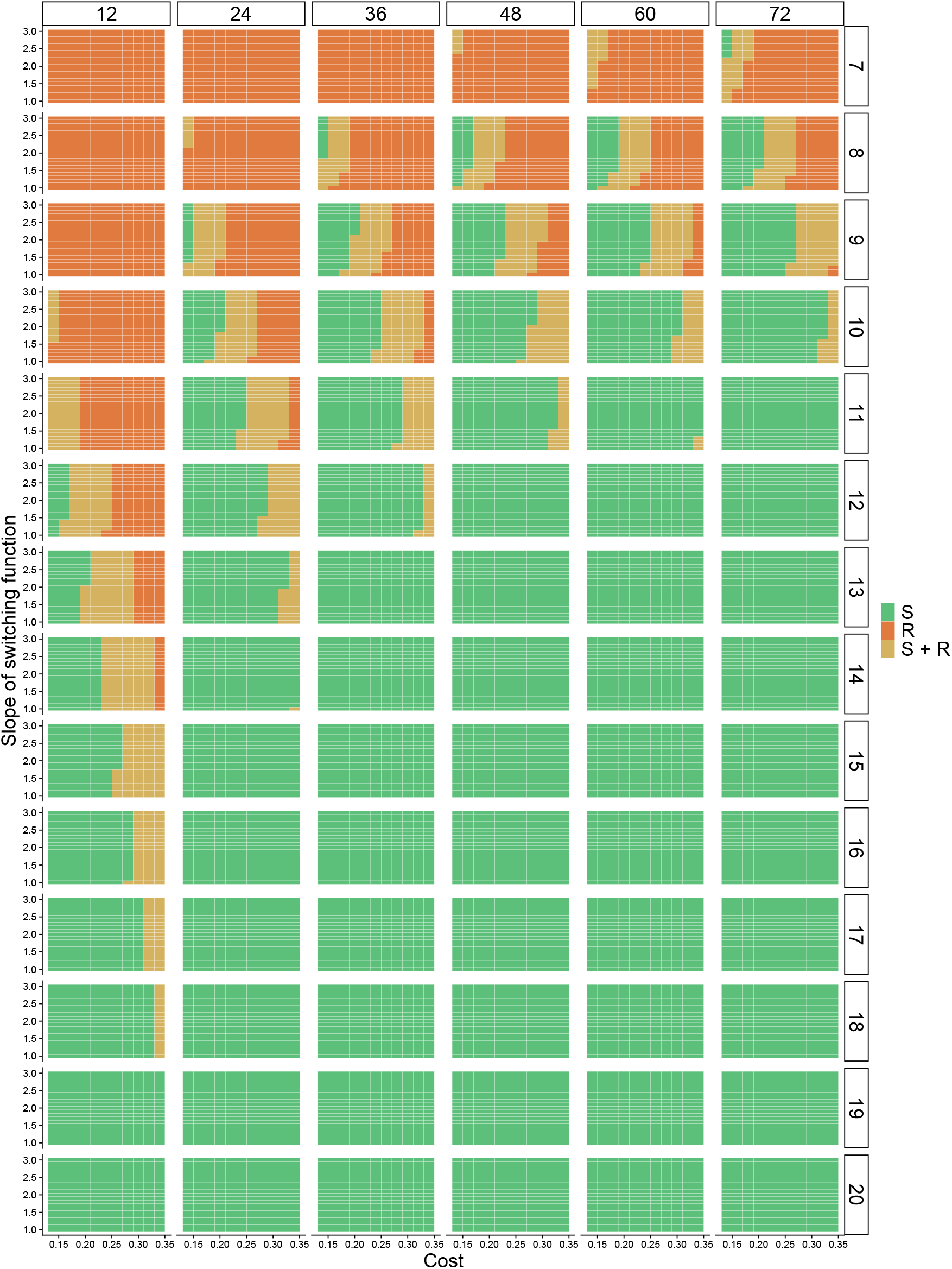
Competition outcomes among the fermenter (F) and switcher (S) across a range of costs (assumed symmetrical between metabolic states), switching function slopes, and resource fluctuation regimes. Columns represent different pulse intervals (inverse of resource supply frequency), and rows represent different pulse sizes (resource input magnitude).

**Figure S6.**
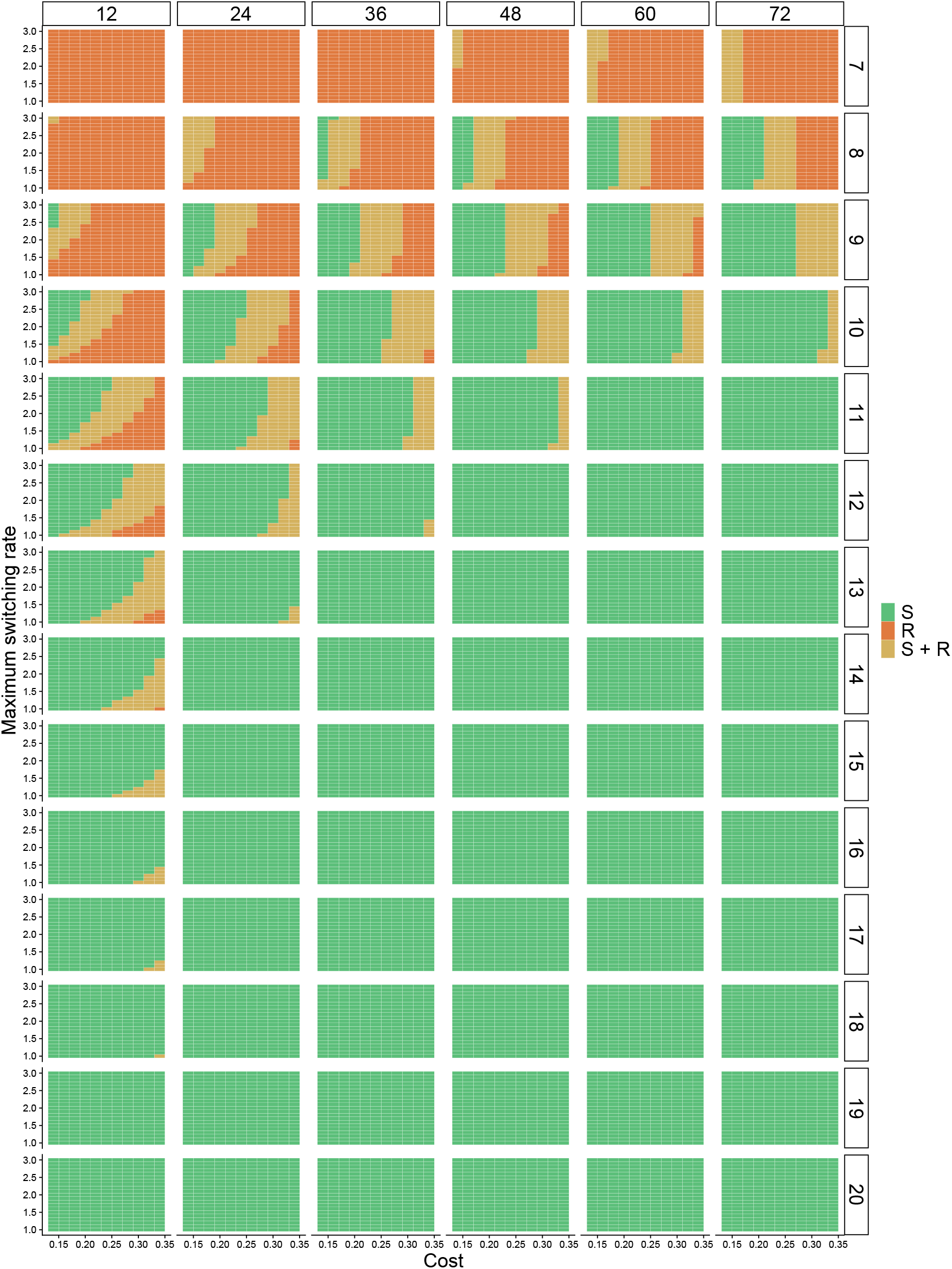
Competition outcomes among the respirer (R) and switcher (S) across a range of costs (assumed symmetrical between metabolic states), the maximum switching rate of the switching function, and resource fluctuation regimes. Columns represent different pulse intervals (inverse of resource supply frequency), and rows represent different pulse sizes (resource input magnitude).

**Figure S7.**
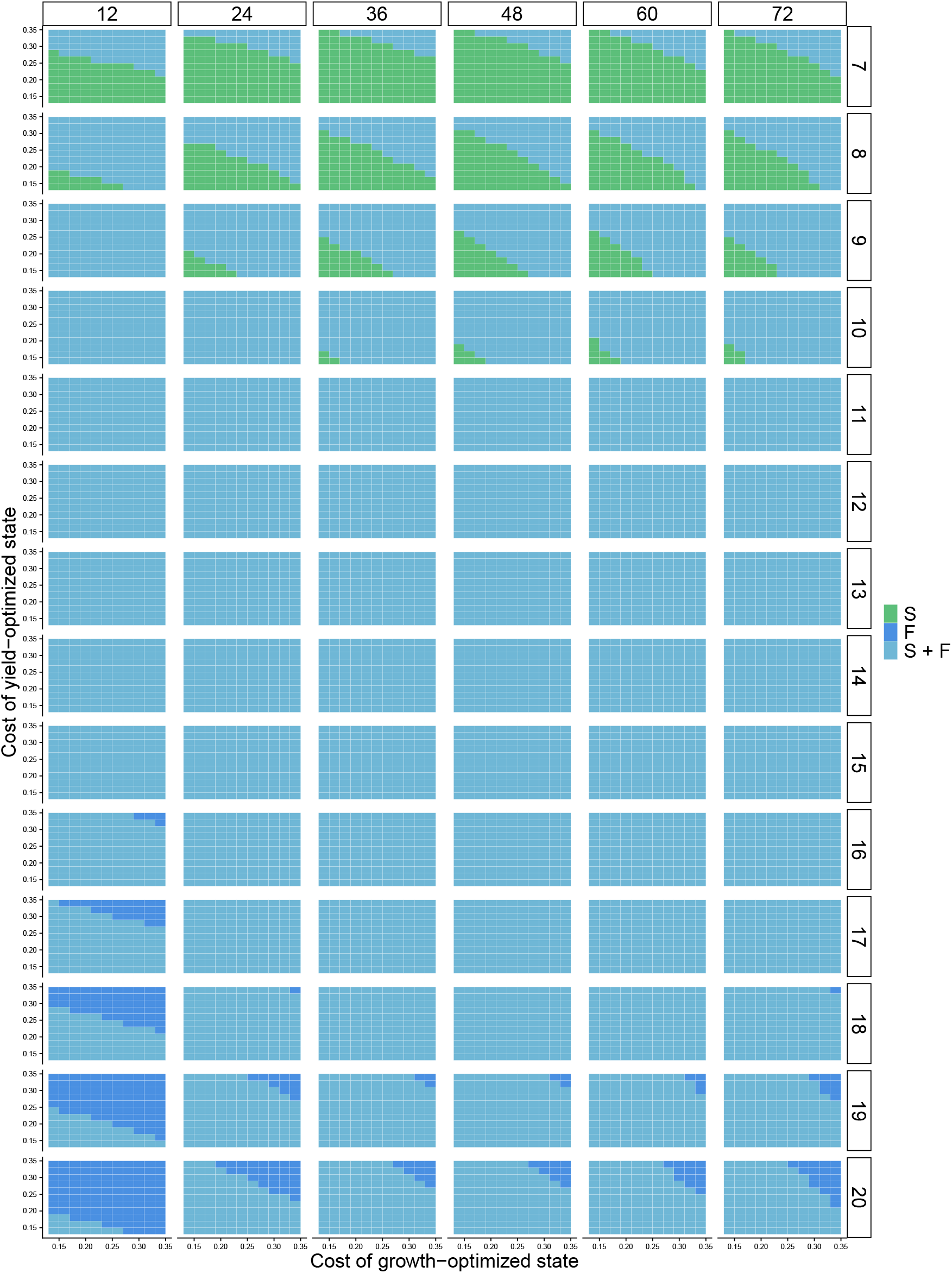
Competition outcomes among the fermenter (F) and switcher (S) across a range of costs and resource fluctuation regimes. Columns represent different pulse intervals (inverse of resource supply frequency), and rows represent different pulse sizes (resource input magnitude).

**Figure S8.**
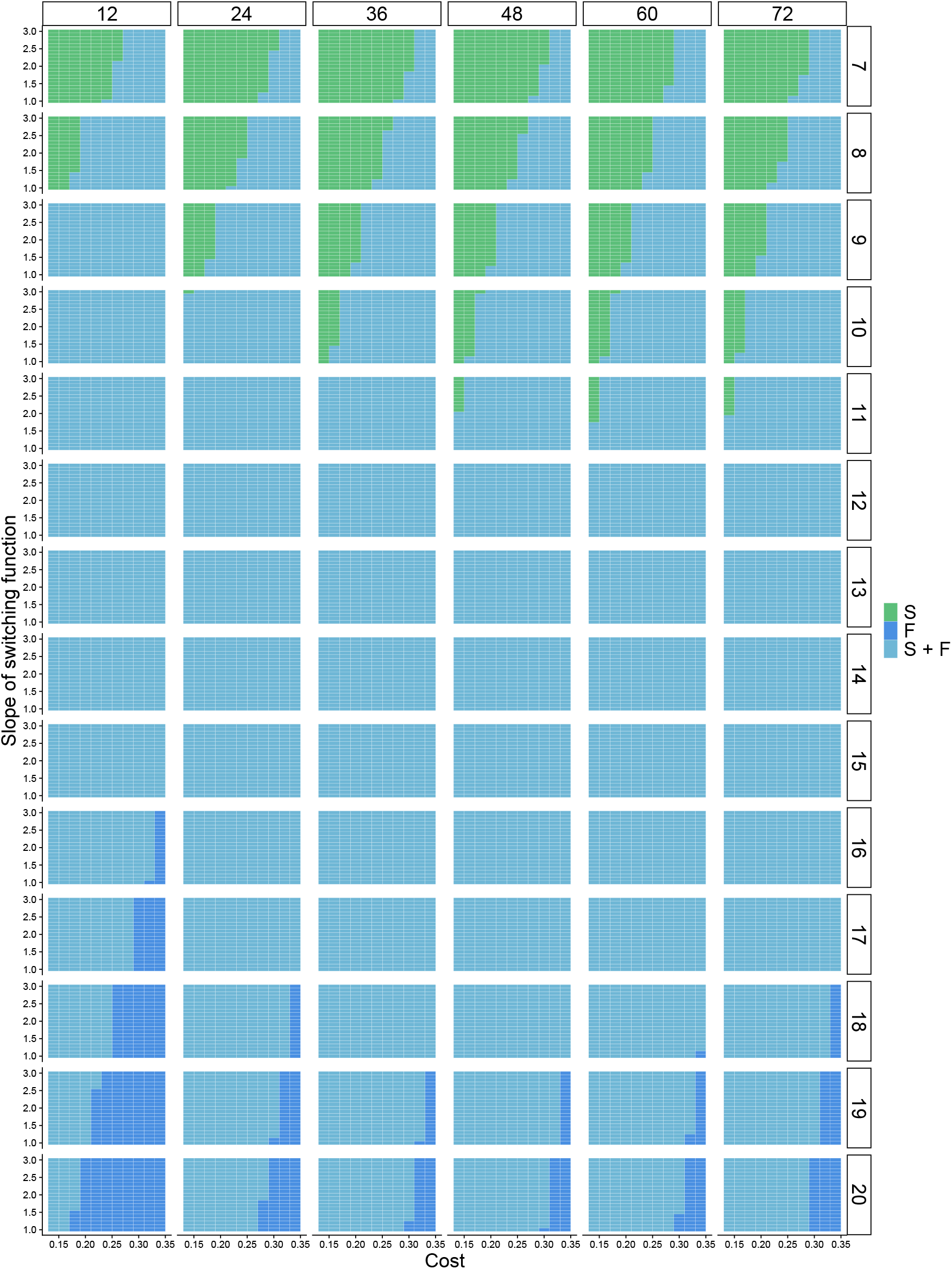
Competition outcomes among the fermenter (F) and switcher (S) across a range of costs (assumed symmetrical between metabolic states), switching function slopes, and resource fluctuation regimes. Columns represent different pulse intervals (inverse of resource supply frequency), and rows represent different pulse sizes (resource input magnitude).

**Figure S9.**
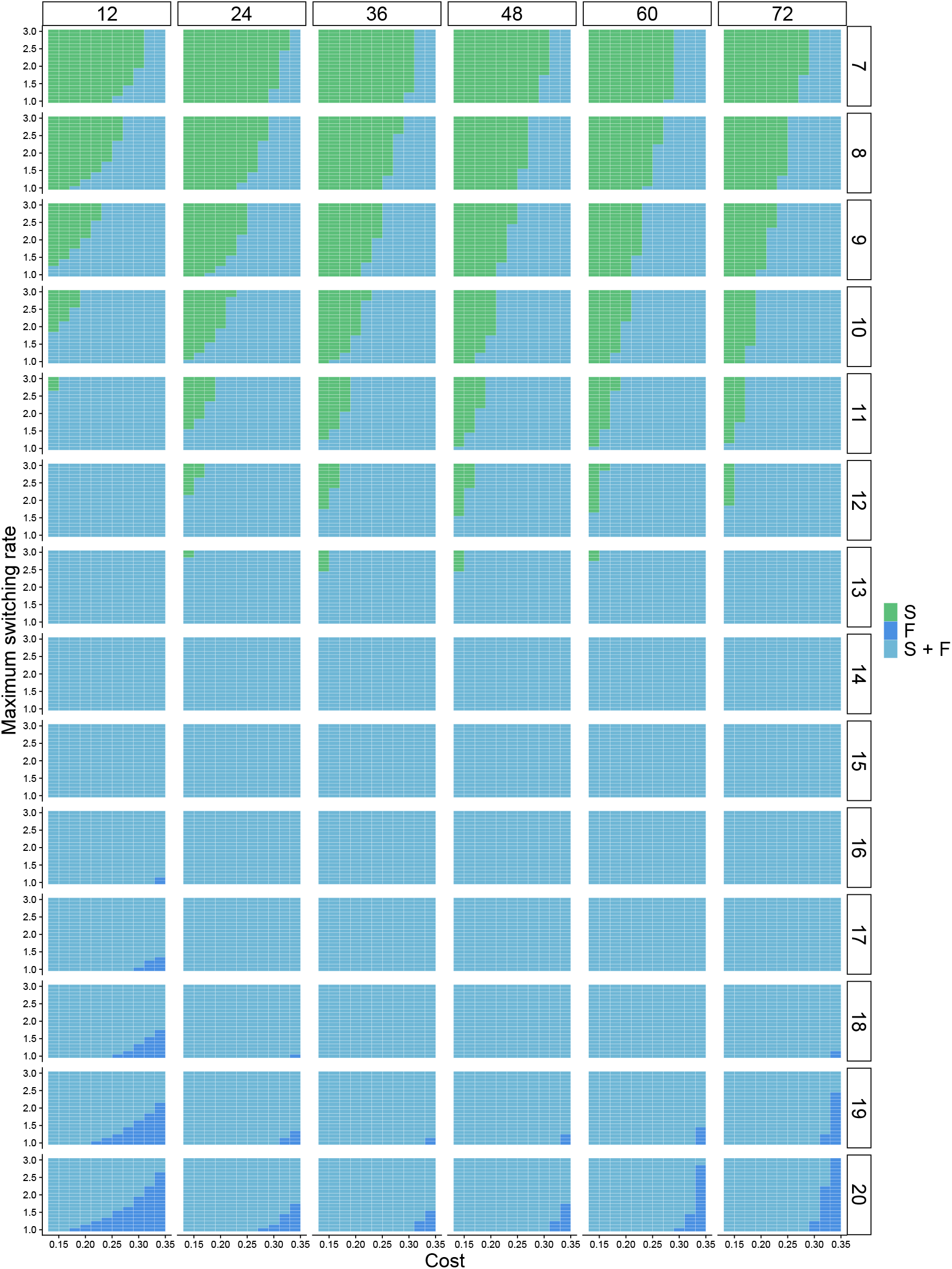
Competition outcomes among the fermenter (F) and switcher (S) across a range of costs (assumed symmetrical between metabolic states), the maximum switching rate of the switching function, and resource fluctuation regimes. Columns represent different pulse intervals (inverse of resource supply frequency), and rows represent different pulse sizes (resource input magnitude).

**Figure S10.**
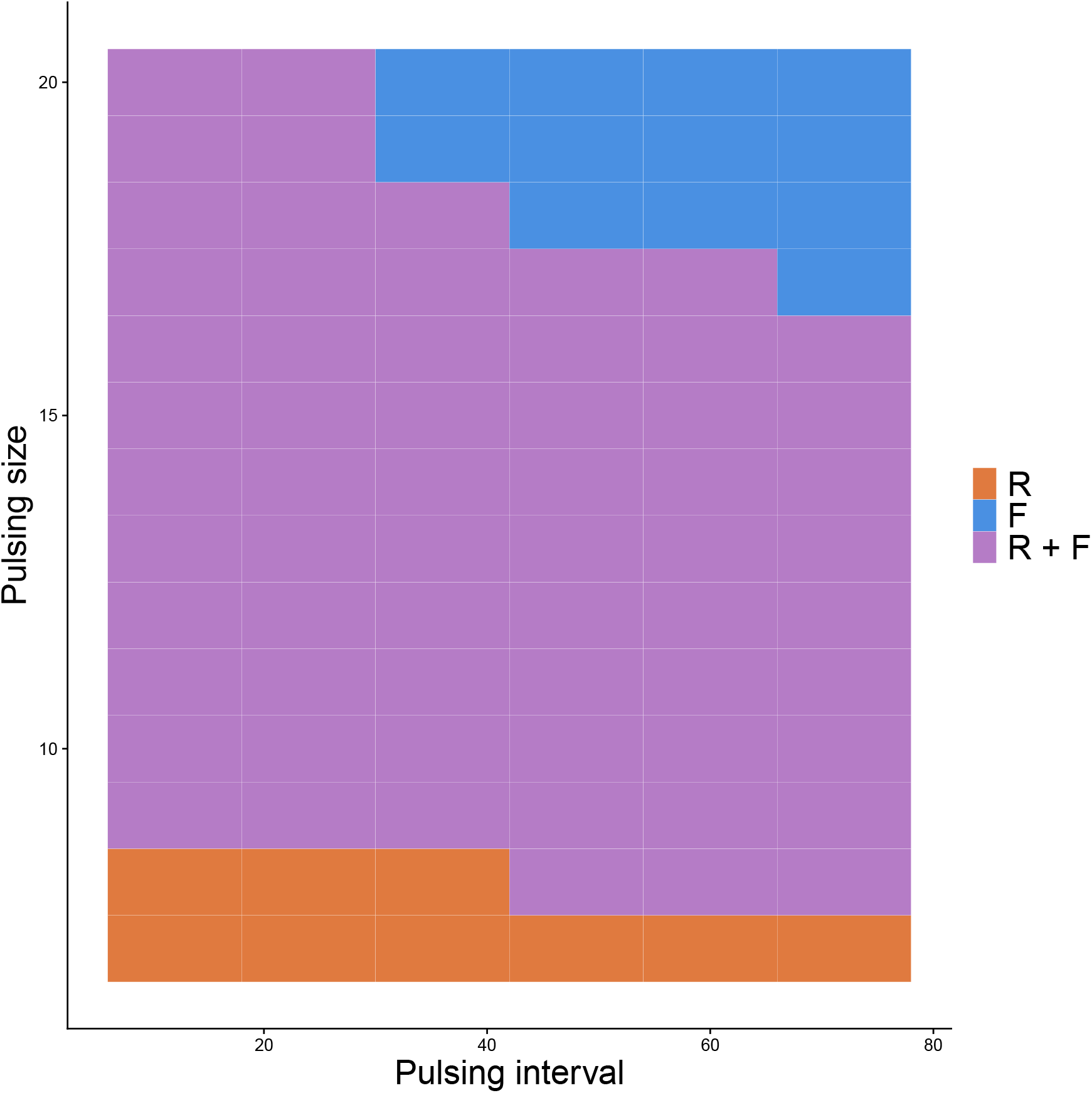
Competition outcomes among the fermenter (F) and respirer (R) across a range of resource fluctuation regimes. Columns represent different pulse intervals (inverse of resource supply frequency), and rows represent different pulse sizes (resource input magnitude).

## References

Acar, M., Mettetal, J.T. & Van Oudenaarden, A. (2008) Stochastic switching as a survival strategy in fluctuating environments. Nature genetics 40, 471–475.

Basan, M., Hui, S., Okano, H., Zhang, Z., Shen, Y., Williamson, J.R. & Hwa, T. (2015) Overflow metabolism in escherichia coli results from efficient proteome allocation. Nature 528, 99–104.

Bauchop, T. & Elsden, S. (1960) The growth of micro-organisms in relation to their energy supply. Microbiology 23, 457–469.

Carlson, R.P., Beck, A.E., Phalak, P., Fields, M.W., Gedeon, T., Hanley, L., Harcombe, W.R., Henson, M.A. & Heys, J.J. (2018) Competitive resource allocation to metabolic pathways contributes to overflow metabolisms and emergent properties in cross-feeding microbial consortia. Biochemical Society Transactions 46, 269–284.

Chang, C.Y., Bajić, D., Vila, J.C., Estrela, S. & Sanchez, A. (2023) Emergent coexistence in multispecies microbial communities. Science 381, 343–348.

Chesson, P. (1994) Multispecies competition in variable environments. Theoretical Population Biology 45, 227–276.

Chesson, P. (2000) Mechanisms of maintenance of species diversity. Annual review of Ecology and Systematics 31, 343–366.

DeLong, J.P., Coblentz, K.E. & Uiterwaal, S.F. (2025) Are type 3 functional responses just statistical apparitions? Ecosphere 16, e70247.

Edwards, K.F., Klausmeier, C.A. & Litchman, E. (2013) A three-way trade-off maintains functional diversity under variable resource supply. The American Naturalist 182, 786–800.

Frank, S.A. (2010) The trade-off between rate and yield in the design of microbial metabolism. Journal of Evolutionary Biology 23, 609–613.

Friedman, J., Higgins, L.M. & Gore, J. (2017) Community structure follows simple assembly rules in microbial microcosms. Nature ecology & evolution 1, 0109.

Gancedo, J.M. (1998) Yeast carbon catabolite repression. Microbiology and molecular biology reviews 62, 334–361.

Grover, J.P. (1997) Resource competition, vol. 19. Springer Science & Business Media.

Kalinkat, G., Rall, B.C., Uiterwaal, S.F. & Uszko, W. (2023) Empirical evidence of type iii functional responses and why it remains rare. Frontiers in Ecology and Evolution 11, 1033818.

Kehe, J., Ortiz, A., Kulesa, A., Gore, J., Blainey, P.C. & Friedman, J. (2021) Positive interactions are common among culturable bacteria. Science advances 7, eabi7159.

Letten, A.D., Yamamichi, M., Richardson, J.A. & Ke, P.J. (2024) Microbial dormancy supports multi-species coexistence under resource fluctuations. Ecology letters 27, e14507.

Levins, R. (1979) Coexistence in a variable environment. The American Naturalist 114, 765–783.

MacLean, R.C. & Gudelj, I. (2006) Resource competition and social conflict in experimental populations of yeast. Nature 441, 498–501.

Millard, P., Enjalbert, B., Uttenweiler-Joseph, S., Portais, J.C. & Létisse, F. (2021) Control and regulation of acetate overflow in escherichia coli. Elife 10, e63661.

Monod, J. (1949) The growth of bacterial cultures. Annual Review of Microbiology 3, 371–394.

Pasteur, L. (1857) Mémoire sur la fermentation alcoolique. Comptes rendus hebdomadaires des séances de l’Académie des sciences 45, 1032–1036.

Pfeiffer, T. & Bonhoeffer, S. (2004) Evolution of cross-feeding in microbial populations. The American Naturalist 163, E126–E135.

Pfeiffer, T., Schuster, S. & Bonhoeffer, S. (2001) Cooperation and competition in the evolution of atp-producing pathways. Science 292, 504–507.

Richardson, J.A., Engelstädter, J. & Letten, A.D. (2025) A unifying principle for multispecies coexistence under resource fluctuations. Proceedings of the National Academy of Sciences 122, e2424996122.

Soetaert, K., Petzoldt, T. & Setzer, R.W. (2010) Solving differential equations in r: package desolve. Journal of statistical software 33, 1–25.

Treves, D.S., Manning, S. & Adams, J. (1998) Repeated evolution of an acetate-crossfeeding polymorphism in long-term populations of escherichia coli. Molecular biology and evolution 15, 789– 797.

Vazquez, A. & Oltvai, Z.N. (2016) Macromolecular crowding explains overflow metabolism in cells. Scientific Reports 6, 31007.

Vemuri, G.N., Altman, E., Sangurdekar, D., Khodursky, A.B. & Eiteman, M. (2006) Overflow metabolism in escherichia coli during steady-state growth: transcriptional regulation and effect of the redox ratio. Applied and environmental microbiology 72, 3653–3661.

Voegele, R.T., Sweet, G.D. & Boos, W. (1993) Glycerol kinase of escherichia coli is activated by interaction with the glycerol facilitator. Journal of bacteriology 175, 1087–1094.

Wanner, B.L., Kodaira, R. & Neidhardt, F.C. (1978) Regulation of lac operon expression: reappraisal of the theory of catabolite repression. Journal of bacteriology 136, 947–954.

Yamamichi, M. & Letten, A.D. (2021) Rapid evolution promotes fluctuation-dependent species coexistence. Ecology Letters 24, 812–818.

Yamamichi, M. & Letten, A.D. (2022) Extending the gleaner–opportunist trade-off. Journal of Animal Ecology 91, 2163–2170, _eprint: https://besjournals.onlinelibrary.wiley.com/doi/pdf/10.1111/1365-2656.13813.

Zhuang, K., Vemuri, G.N. & Mahadevan, R. (2011) Economics of membrane occupancy and respirofermentation. Molecular systems biology 7, MSB201134.

